# Modeling whole brain Electroencephalogram (EEG) in a spatially organized oscillatory neural network

**DOI:** 10.1101/2023.07.16.549247

**Authors:** Sayan Ghosh, Dipayan Biswas, Sujith Vijayan, V. Srinivasa Chakravarthy

**Author notes:** these authors contributed equally to this work.

## Abstract

In this study, we model high-dimensional Electroencephalogram signals in sleep stages, using a general trainable network of Hopf oscillators. The proposed architecture has two components: a layer of oscillators with lateral connections, and a complex valued feedforward network with and without a hidden layer. The output of the Hopf oscillators, whose dynamics is described in the complex domain, is fed as input to the feedforward network and the output predicts the EEG signals. The proposed network is trained in two stages: in the 1st stage, intrinsic frequencies of the oscillators and the lateral connections are trained whereas, in the 2nd stage, the complex-valued feed-forward network is trained. Reconstruction error obtained when there is a hidden layer in the feedforward network is an order of magnitude smaller than that obtained without a hidden layer. Also, it has been shown that during testing our model is able to generate EEG signals, whose spectral properties accurately describe the corresponding target signals. In the first, the oscillators do not have any spatial localization, whereas in the second the oscillators are spatially distributed in a spherical shell geometry. The model described can be interpreted as a stepping stone towards a large-scale model of brain dynamics.

## Introduction

Using extracranial electrodes, EEG measures the extracellular ionic current produced by a graded postsynaptic potential of vertically oriented pyramidal neurons in the III, V, and VI cortical layers [1]. The electrical dipole field created by the soma and apical dendrites of pyramidal neurons is propagated through layers of the cortex, cerebrospinal fluid (CSF), skull, and scalp via volume conduction [2] and recordable at the scalp site [3]. EEG technology has diverse applications including characterizing brain dynamics in early stages of Parkinson’s disease (PD) [4], epileptic seizure detection [5], motor imagery and movement classification in brain-computer interfaces [6], emotion classification [7], etc. Sleep is a complex naturally recurring dynamic process that occurs periodically in most animals [8]. There are three key physiological mechanisms or rhythms that regulate sleep, viz., circadian rhythm, homeostasis, and ultradian rhythm [9-10]. Polysomnogram [11], which jointly measures brain electrical activity (EEG), muscle activity (Electromyogram-EMG), eye movement (Electrooculogram-EOG), and heart rate (ECG) is a standard method of recording sleep activity. Sleep stages can be broadly categorized into 5 stages: wake, non-rapid eye movement N1 (NREM N1), non-rapid eye movement N2 (NREM N2), non-rapid eye movement N3 (NREM N3), and rapid eye movement (REM) sleep [12]. Sleep facilitates important neural and physiological functions including memory consolidation [13], emotion control [14]. In the past decade, several efforts have been made to model the brain as a nonlinear dynamical system and describe brain dynamics using complex nonlinear dynamical networks [15]. A phenomenological model comprising van der Pol–Duffing double oscillator networks was used to model EEG signals from healthy controls as well as Alzheimer disease patients [16]. The model results compare favourably with experimental results in terms of time series, power spectrum, and Shannon entropy [16-18]. Another model accounted for the generation of EEG ictal patterns from the temporal lobe using coupled Duffing-van der pol oscillators [19]. By an analysis of EEG time series, the presence of low dimensional chaos in NREM N1 and REM sleep stages was described by Babloyantz et al. [20]. Efforts have been made to use stochastic limit cycle oscillators to model EEG data from healthy subjects [21-22]. Despite these efforts, the link between this mesoscopic EEG activity and the dynamics underlying neuronal circuits is still not fully unraveled. The weighted mean potential of a weakly-coupled, local cluster of Hindmarsh-Rose (HR) neurons collectively shows near-synchronization behavior and can optimally reconstruct epileptic EEG time series [23]. A similar line of work was also proposed by Phuong and colleagues, in which networks of Hindmarsh-Rose (HR) neurons and Kuramato oscillators were used to reconstruct EEG data in healthy and epileptic conditions [24]. There is a growing interest in modeling large scale brain activity using networks of nonlinear oscillators. A notable example of this kind is The Virtual Brain (TVB) framework, which uses large oscillatory networks to model various manifestations of functional brain dynamics like EEG, functional Magnetic Resonance Imaging (fMRI), and, Magnetoencephalogram (MEG) [25]. Al-Hossenat et al. proposed modeling of slow wave activity (delta and theta) of EEG from eight different regions using Jansen and Rit (JR) neural mass model and anatomical connectivity using the TVB framework [26]. In another modeling study, using a similar kind of neural mass model, alpha wave activity was reproduced from four different brain regions [27]. In this paper, we propose a network of Hopf oscillators described in the complex domain and show how the network can be trained to model high-dimensional EEG data in waking and sleep stages. In an earlier work we showed how to achieve a stable phase relationship between a pair of oscillators, with arbitrarily different frequencies, using a special form of coupling known as power coupling [28]. With power coupling the difficulties that arise in a pair of coupled oscillators, depicted by Arnold tongues, seem to be overcome effectively. It was shown how networks of such coupled oscillator systems can be trained to learn a small number of EEG channels. In the present study, we add a hidden layer of sigmoidal neurons, and geometrically constrain the network, so as to learn high-dimensional, “whole brain” EEG signals accurately. The outline of the paper is as follows. This article begins with an account of sleep EEG recording followed by preprocessing methodology. The two stages of training of the proposed network are described in ’1st stage of training’ and ’2nd stage of training’ section, respectively. Section ’Insertion of hidden layer’ describes the deep oscillator network, which is a combination of oscillatory layer and a feedforward network. Section ’Generation of EEG Data’ describes the generation of EEG data using Hopf oscillator model. Section ’spatial distribution of oscillator’ shows how oscillators are distributed on a rectangular grid or spherical shell-geometry. Sections ’Reconstruction without hidden layer’ and ’reconstruction with hidden layer’ illustrate the model performance with two kinds of feedforward network – one with and one without a hidden layer. A discussion of the work is presented in the last section.

## methods

### EEG Recording

Polysomnogram (PSG) datasets comprising 56 EEG electrodes, 2 EoG electrodes and four EMG electrodes, were recorded from two healthy subjects (full night, 8hr) at the School of Neuroscience, Virginia Tech, USA. During data collection, all necessary instructions, such as those regarding caffeine and alcohol use restrictions, were adhered to. The different stages of sleep are scored according to American Academy of Sleep Medicine (AASM) rules by two sleep experts [29]. A night’s sleep consists of periods of rapid eye movement (REM) sleep and periods of non-rapid eye movement (NREM) sleep; the latter consists of three stages, NREM1 (N1), NREM2 (N2), and NREM3 (N3), also known as slow wave sleep. NREM N1 is often the stage when a transition occurs between waking and sleep; the awake state is characterised by low amplitude and relatively high frequency waves. NREM N1 occurs for 3–8% of the duration of total full night sleep and is dominated by theta waves (4-7 Hz) [8]. NREM N2 is defined by sleep spindles (11-16 Hz) and K-complexes. NREM N3 is also called slow-wave sleep as there is prominent activity in the 0.1 to 4 Hz range. REM sleep is somewhat similar to the wake stage, which occurs more frequently late at night and occupies 20% of total sleep [9]. REM sleep is marked by muscle atonia and conjugate eye movements. These datasets are in European Data Format (.EDF) format. For further analysis, we converted these datasets into .mat format using the EEGLAB plugin in Matlab, and extracted 10 sec, and 30 sec long chunks from each of the 56 channels of EEG data.

### Pre-processing

EEG is inherently a noisy signal influenced by non-neural factors e.g., (muscle movement measured by Electromyogram (EMG), eye movement measured by Electrooculogram (EOG), and Electrocardiogram (ECG)), as well as equipment noise (power line interference (50/60 Hz), impedance fluctuation and, cable movements [29]. Here we applied bandpass filtering from 0.1 Hz to 20 Hz using Butterworth approximation with filter order 2 and stopband ripple 3 dB and passband ripple 40 dB. In addition, EEG data is normalized to remove DC noise. The sampling frequency of the system is 500 Hz.

### A network of Neural Oscillators

For our current purpose of modelling multi-dimensional EEG signals, we use an enhanced version of a network of neural oscillators described in [28]. The original model of [28] consists of a layer of Hopf oscillators with lateral coupling connections, and an output layer which is directly connected by a single weight stage to the oscillator layer. Dynamics of the Hopf oscillators was described in the complex domain, coupled using a special form of coupling known as power coupling. The layer of oscillators is connected to the output layer using all-to-all linear forward weights. Thus, the given time series is modelled as a linear sum of the outputs of the layer of oscillators. In the present model, a hidden layer of sigmoid neurons is inserted between the oscillatory layer and the output, which immensely reduces the fitting error. The original network of [28] has two components: the input oscillatory layer consisting of a network of coupled Hopf oscillators, and a feedforward linear weight stage that maps the outputs of the oscillators onto the network’s output node(s). We use the Hopf oscillator in the supercritical regime where the oscillator exhibits a stable limit cycle. In the previous study, we introduced ’power coupling’, which shows a way of achieving a constant normalized phase difference between a pair of coupled Hopf oscillators with arbitrary intrinsic frequencies [28]. The dynamics of the oscillatory layer are described by eqns. (1-5) below.

### The Single Hopf oscillator

Complex domain representation of single Hopf oscillator is described as:

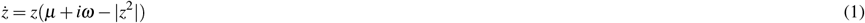

where z is a state variable –

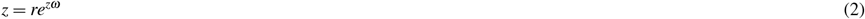

The dynamics of N coupled Hopf oscillators without external input can be described as:

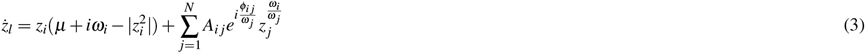

The polar coordinate representation of eqn. 3 is

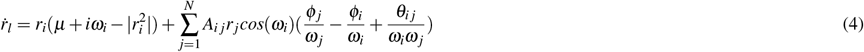

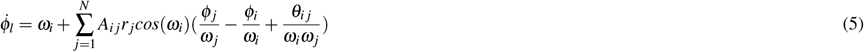

where r and *ϕ* are the state variables, 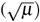, (*μ*>0) is the amplitude of oscillation and *β* is a bifurcation parameter. *μ* = 1, *β* = 20. *A*_*i j*_, is the magnitude of complex coupling coefficient, (*A*_*i j*_ << 1). *θ*_*i j*_ is angle of complex coupling coefficient, *ϕ*_*i*_ and *ω*_*i*_ and are the ith oscillator’s phase and natural intrinsic frequency respectively.

The network described above is trained in two phases: in stage 1, the intrinsic frequencies, *ω*_*i*_, of the oscillators and the coupling weights among the oscillators 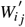 in the oscillatory layer are trained; in stage 2, the feedforward linear weights between the oscillatory layer and the network output are trained.

### 1st stage of training

Since the aim of the 1st stage of training is to train the intrinsic frequencies, *ω*_*i*_, these frequencies are initialized by sampling from a uniform random distribution over the interval [0, 20] Hz. The modified network dynamics is described in eqn. 4, where error signal e(t) drives each oscillator.

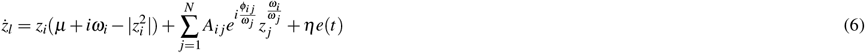

The teaching signal used for training is denoted by, D(t), which is an EEG signal of a finite duration. The power coupling weight, *W*_*i j*_, in eqn. (7), is the complex lateral connection, *θ*_*i j*_ is the angle of lateral connection and *A*_*i j*_ is the magnitude of the lateral connection between the 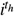 and 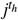 oscillators, and *ω*_*i*_, is the intrinsic frequency of the i’th oscillator. The oscillator activations are summed using feedforward weights,*α*_*i*_, which, in this stage of training, are taken to be small real numbers 0.2 (i.e. *α*_*i*_=0.2 for all i) and lateral connection 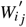 is initialized with complex numbers according to eqn. (7). S(t) is the predicted or network reconstructed signal shown in eqn. (11). The whole network is driven by an error signal e(t) which is the difference between network predicted signal and the desired signal eqn. (9). The training of intrinsic frequency, *ω*_*i*_, is described in eqn. (8) where *η*_*ω*_ is the learning rate, e(t) is the error signal and *ϕ*_*j*_ is the oscillator phase. Training of the real feedforward weights is done by modified delta rule, eqn. (10), where *η*_*α*_ is the learning rate of feed forward weight update. Basically, the network performs some sort of Fourier decomposition of the target signal, and each oscillator is used to learn the frequency component closest to its own intrinsic frequency present in the signal. In the 1st stage, the intrinsic frequency, the angle of power coupling weight, and amplitude of the signal (real feedforward weight) are trained. Lateral connections are trained by a complex-valued Hebbian rule (eqn. 12). Hebbian learning of complex power coupling is shown in equn. (12) Weight of power coupling can be written as

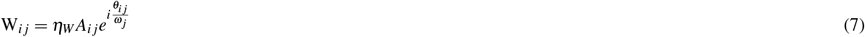

where *τ*_*w*_ is the time constant.

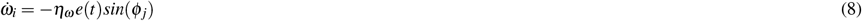

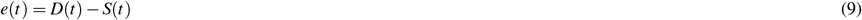

Where D(t) is the teaching EEG time series and S(t) is the network predicted time series.

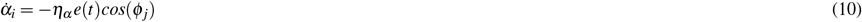

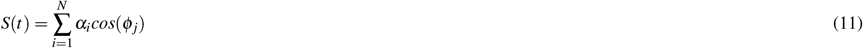

Hebbian learning of complex power coupling is shown in equn(10).

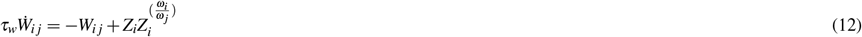

where *τ*_*w*_ is the time constant.

### 2nd stage of training

In the 2nd stage of training, the oscillatory network with learned intrinsic frequencies and lateral connections of the oscillatory layer from the previous stage was used as a starting point. (Note that the oscillatory layer with trained parameters may be compared to a reservoir of reservoir computing [28].) But the main difference is that, in this stage the feedforward weights are allowed to be complex (they were real in 1st stage training) and trained once again by supervised batch mode learning rule. *K*_*i j*_ and *ζ*_*i j*_ are the magnitude and angle of complex feed forward weights updated according to eqns. (3,4) in supplementary file. The complex feedforward weights and network ouput can be defined as :

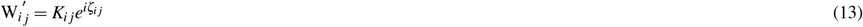

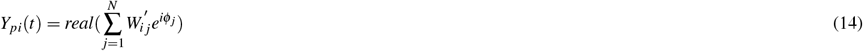

### Insertion of Hidden Layer

As we will see subsequently in the result section, despite the theoretical advantages, the model described above does not yield satisfactory approximations of the empirical EEG signals. In order to improve the approximation performance, we insert a hidden layer of sigmoidal neurons between the oscillatory layer and the output. In the new version of the model with the hidden layer, the intrinsic frequencies of the oscillators and their lateral connections are trained using the learning mechanisms of the 1st stage of training described above (eqns. 4-12).

In the 2nd stage of training, the oscillator activation is presented via a complex-valued, fully connected weight stage to a hidden layer with a complex-valued tanh activation. Similarly, the hidden neurons and the output layer are also connected by a complex-valued, fully-connected, feedforward weight stage. In this module initially EEG signal is trained during stage 1 with real feedforward weights (where the oscillator’s’ intrinsic frequencies are trained) and in stage 2 has two substages where linear complex weights are multiplied with oscillator’s activation followed by a nonlinear stage.

### Generation of EEG Data

So far, using supervised training, we have only reconstructed the data. To validate our model, we have to generate the output of the model without the network being driven by training signal. The network can generate a 30-second EEG signal without any external input. During generation, the initial phases of oscillators are initialized randomly but the natural intrinsic frequency of oscillators, all feedforward weights (oscillatory layer to hidden layer weights and hidden layer to output layer weights) are adopted from the trained network. Here r and dynamics (supplementary file: eqn. (27-28)) are derived from complex domain (eqn. 4) by transforming to polar coordinates. The time series of newly generated EEG data may not match the desired EEG signal beyond the training duration,but the spectral features of the generated signal must be similar to the testing duration of the raw EEG data. The spectral features of the generated EEG segments are compared over the test duration (the next 30 seconds after the training duration). Note that we extracted the train data in such a way that the next 30 sec data (test data) also belongs to the same sleep stage as that of the training duration.

### Spatial Distribution of Oscillator

Currently, we will try to model the cortical sources using oscillators. To this end, we consider two spatial distributions of the oscillators within a “cortical layer” which is modeled in two ways: 1) a spherical shell 2) a rectangular grid (supplementary file). Electrodes are placed on the top of cortical layer, inside another layer named the ‘electrode layer’. Real-world 10-20 electrode geometry is introduced in the ‘spherical shell’ case.

Next, we specify which oscillators in the cortical layer contribute to which electrodes in the electrode layer. This is done by a simple nearest neighbor criterion: only the oscillators that lie within a threshold distance *ε*_1_ from a given electrode contribute to that electrode, as defined in eqn. (15).

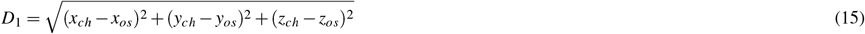

where *D*_1_ is the distance from a given electrode in the electrode layer to a given oscillator in the cortical layer,. Where *D*_1_ is the distance from electrode layer to cortical layer. There is also another layer named ‘hidden layer’ in between the cortical layer and the electrode layer. No specific spatial location is specified for the neurons in the hidden layer. Here basically the model architecture followed was similar to that described in Section II .F, fig. 2 but with one important difference: a separate hidden layer of neurons is introduced for every electrode. In the schematic shown in fig.3, two different hidden layers are depicted corresponding to two distinct electrodes. The corresponding oscillators are also shown. Note that though the hidden layers are not shared between two electrodes, the corresponding oscillators can be partially shared (fig. 3).

**Figure 1.**
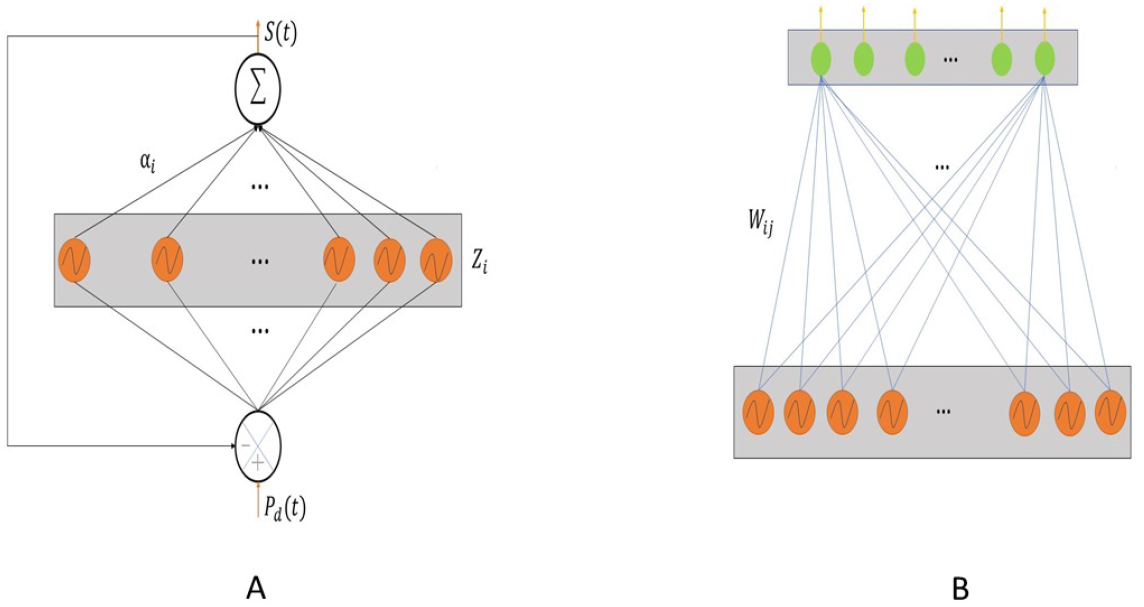
A)- Network architecture of 1st phase of training; B)- Network architecture of 2nd phase of training.

**Figure 2.**
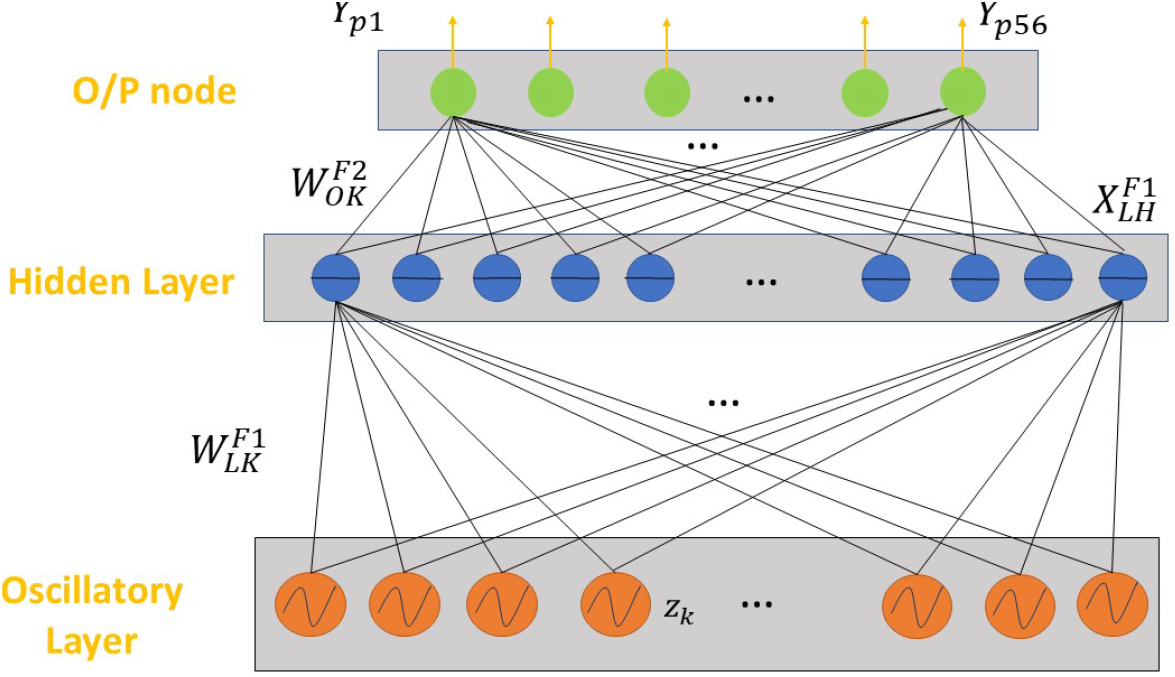
Inserting the hidden Layer between the oscillatory layer and the output layer.

**Figure 3.**
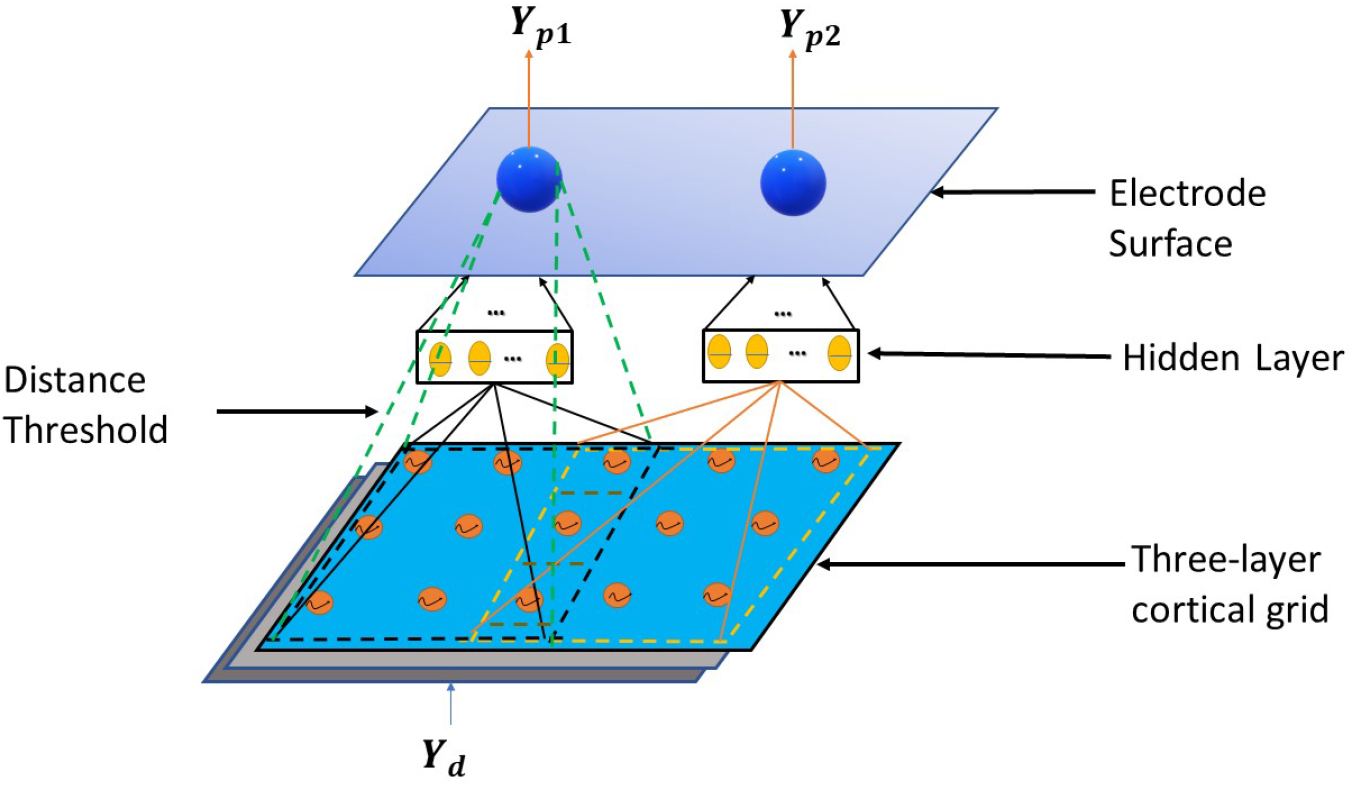
The connections between the oscillators in the cortical layer, the hidden layer and the electrodes in the electrode layer. The dashed green lines indicate the network associated with the electrode on the left in the electrode layer.

The thresholding process mentioned in the above determines electrodes and their corresponding oscillators. We use another threshold which determines the lateral connections among the oscillators. Here we use spatial location of the oscillators in the cortical layer to specify local connectivity using another distance-based threshold (ζ_2_) derived in eqn. (16). Long-range connections among the oscillators are not used in this study.

Consider, 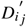 is the distance among oscillators. A pair of oscillators whose mutual distance exceeds the distance threshold limit *ζ*_2_, are not connected. 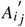 represents the magnitude of the complex-valued connection, 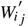, between i’th and j’th vity among oscillators. The magnitude of 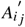 is set to 0.001. Only those oscillator pairs are connected and trained which are within the threshold distance (*ζ*_2_) of each other i.e. ith and jth oscillator are connected only if 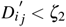 where,

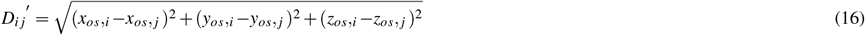

if,
else,

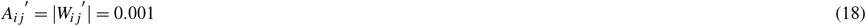

Note that the angle of the coupling connections, 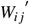, is calculated by Hebbian learning as per eqn. 10.

### Spherical shell

In this section, we describe a model in which the cortical layer is modeled as a spherical shell. Likewise, in reality, EEG electrodes also are not confined to a planar surface. We currently place the electrodes on a spherical surface on top of the spherical cortical layer. We extracted the precise electrode locations from EEGLab [30]. In EEGLab, we used the four-shell dip fit spherical model of Brain Electrical Source Analysis (BESA) to find the individual electrode locations. In the four shell spherical model inner sphere represents the brain and other three outer shells represent CSF(cerebro spinal fluid), skull and scalp. Based on the position and radius of electrode layer we create a spherical shell. A set of 8 electrodes are shown on the cortical layer in fig. (4). Similar to the case of the rectangular grid, in this case too, there is a hidden layer between the cortical layer, which consists of oscillators, and the electrode layer.

**Figure 4.**
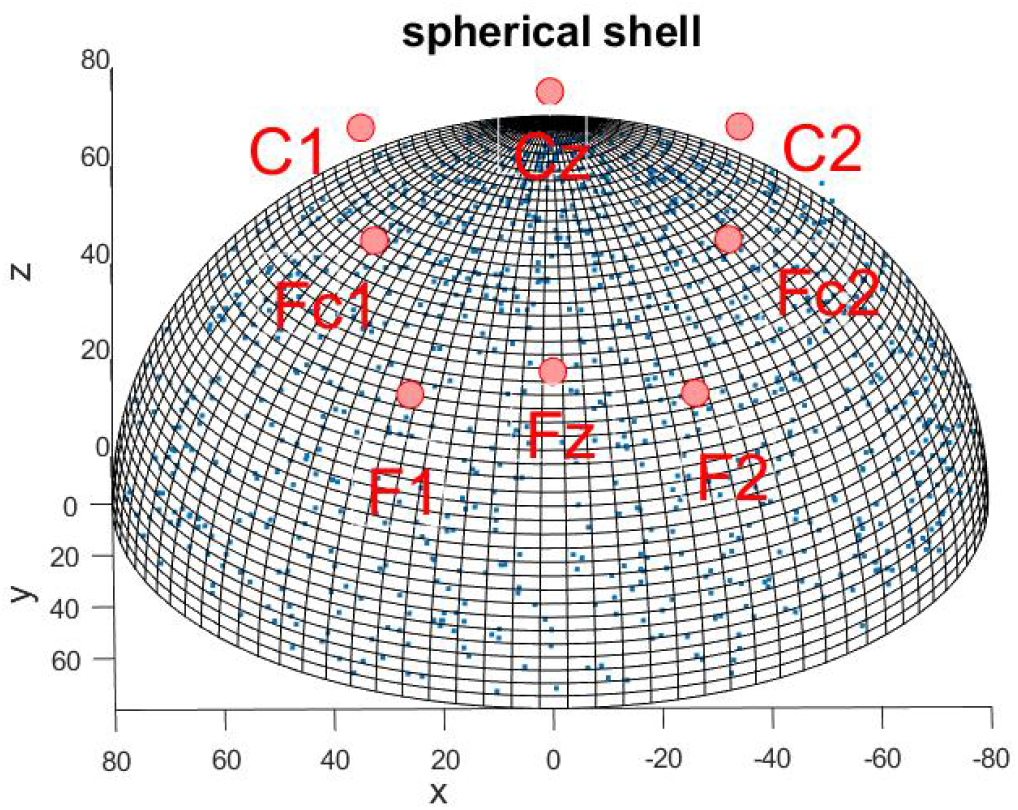
Spherical shell geometry within which the electrodes are placed.

**Figure 5.**
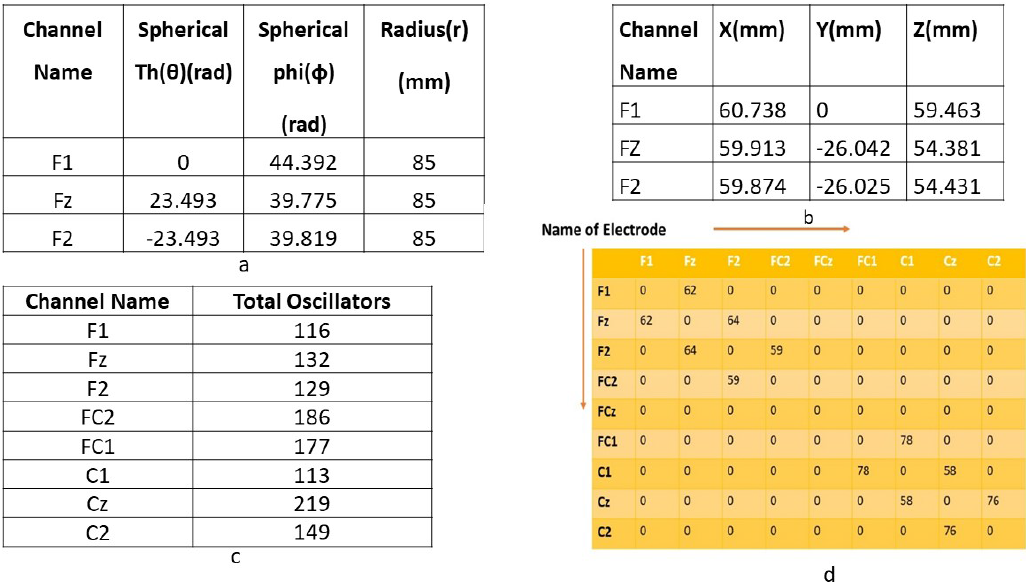
a) Sample location in spherical coordinate system froorm EEGLAB; b)- Sample location in Cartesian coordinate system fromform EEGLAB; c)- The number of oscillators allotted to each electrode; d)- Pair of electrodes and their shared oscillators.

**Figure 6.**
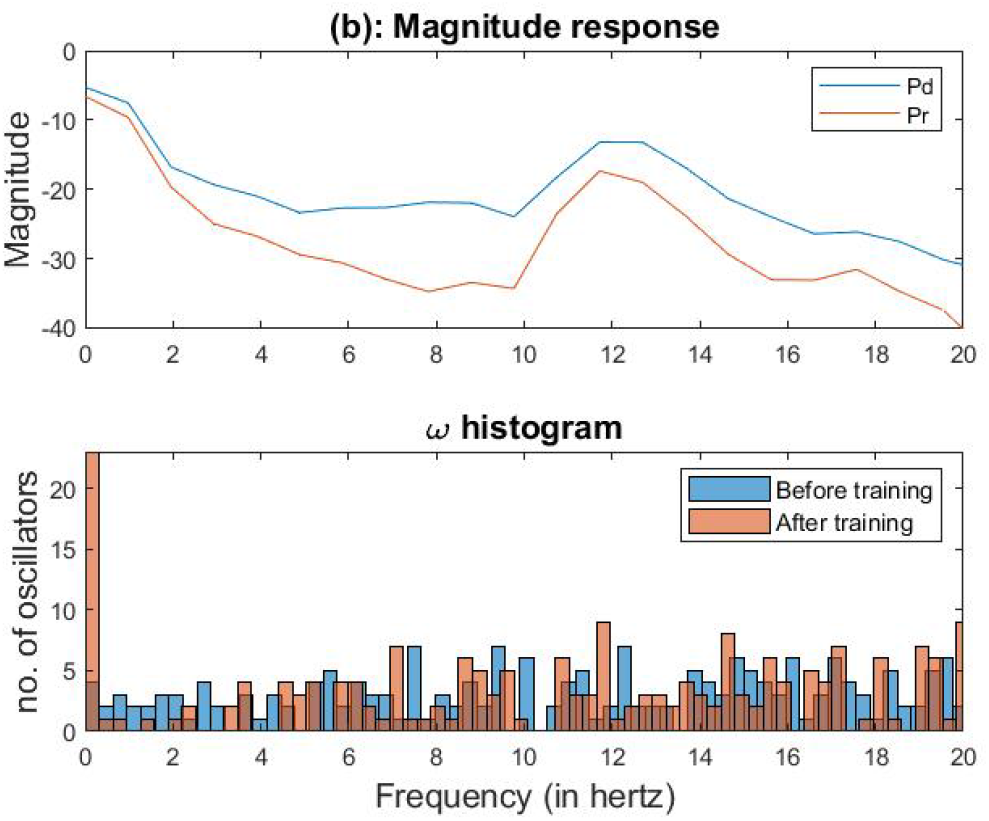
learning the intrinsic frequencies.

## Results

In this section, we describe the performance of the models described in the previous section on modelling high-dimensional, whole brain EEG data (56 electrodes). In order to model a large number of electrodes, as well as essential frequency band (0.1 to 20 Hz), a large number (N=200) of oscillators are used. Although the model is trained on the original EEG time series, the model performance is better demonstrated in spectral terms, rather than in the time domain. In order to depict the signal spectrum, instead of using normal FFT, we use average periodogram method known as the “Welch method” [31]. We use the Welch method with a Hamming window of size 1 sec and 50% overlap throughout the paper.

### Reconstruction without hidden layer-1st stage of training

Following the method described in Section ’1st stage of training’., we show how the intrinsic frequency of the oscillators adapts to the nearby frequency components present in the desired signal. The intrinsic frequencies of the oscillators, *ω*_*i*_, are initialized by drawing from a uniform distribution over the interval [0, 20] Hz. The real feedforward weight (*α*_*i*_) which connects *i*^*t*^*h* oscillator to the single output node is uniformly initialized with a small real number (=0.2). The complex-valued lateral connections, 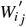, are initialized according to eqn. 7. Note that in this stage, we do not use the hidden layer in the feedforward stage. Training is performed on EEG chunks of duration 10 sec and 30sec (supplimentary file) are separately used for training. Frequency learning rate is *η*_*ω*_ =0.0001 (eqn. 8), amplitude learning rate *η*_*α*_ = 0. 0001 (eqn. 10), and the learning rate for the coefficient of lateral connection weight, which determines the magnitude of oscillator-to-oscillator connections (A), (eqn. 7), is*η*_*w*_ = 0.001

### 2nd stage of training

Following the method described in Section Method. ’2nd stage of training’, in this stage, we use the learned intrinsic frequencies and lateral connections from the 1st stage of training, while amplitude and phases of the complex feed-forward weight (*W*_*i j*_) are trained further. The model-generated signal is computed according to eqn. (14). Although the equation shows a single electrode signal, by using a matrix of feedforward weights we can reconstruct any number of electrode signalss. Therefore, the predicted signals (fig. 7C) from the 2nd stage look better than the 1st stage (fig. 7 A). Whereas in the 1st stage we use real feedforward weights, in the 2nd stage we use complex feedforward weights: this is the only difference between the two stages of training. The performance of the network is evaluated in terms of the Root Means Square Error (RMSE) between the predicted and desired signals for all 56 channels. Reconstruction using 30sec EEG Data is shown in supplementary file (section 5).

**Figure 7.**
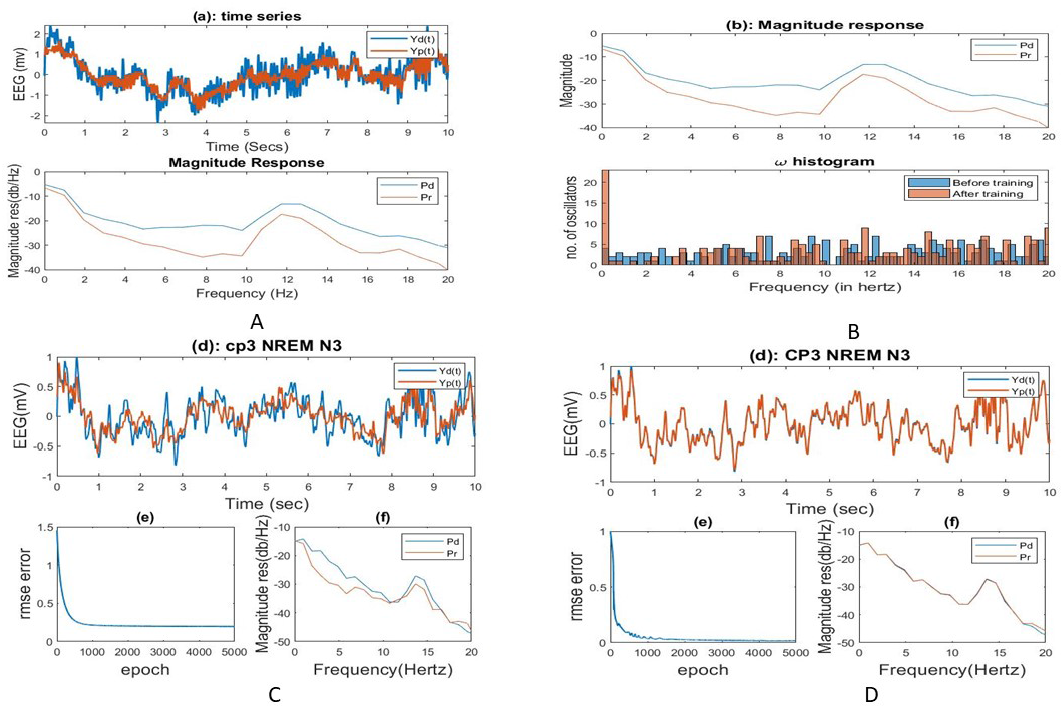
A)- Time series reconstruction after 1st phase of training; B). learning of intrinsic frequency C). time series reconstruction of CP3 (without hidden layer) D). Time series reconstruction after inserting the hidden layer.

### Reconstruction with hidden layer

In the previous subsection, we observed that the reconstruction error is poor when there is no hidden layer. To improve the model performance, following the method described in Section Method. ’insertion of hidden layer’., we inserted a hidden layer of 100 sigmoidal neurons between the oscillatory layer and the output layer in the 2nd stage of training. This modification greatly improved performance, with the RMSE values dropping (by an order of magnitude). Comparisons between the cases of ‘without hidden layer’ and ‘with hidden layer’ model prediction for all the five stages of sleep are shown in a bar plot (fig. 8). Then we compared those two Phase locking value(PLV) matrices using linear correlation and observed the similarity among ‘model predicted’ and ‘Experimental’ data in terms of functional connectivity. And we obtained correlation values 0.5517 and 0.9903 in case of ‘without hidden layer’ and ‘with hidden layer’ respectively (supplementary file: section 4).

**Figure 8.**
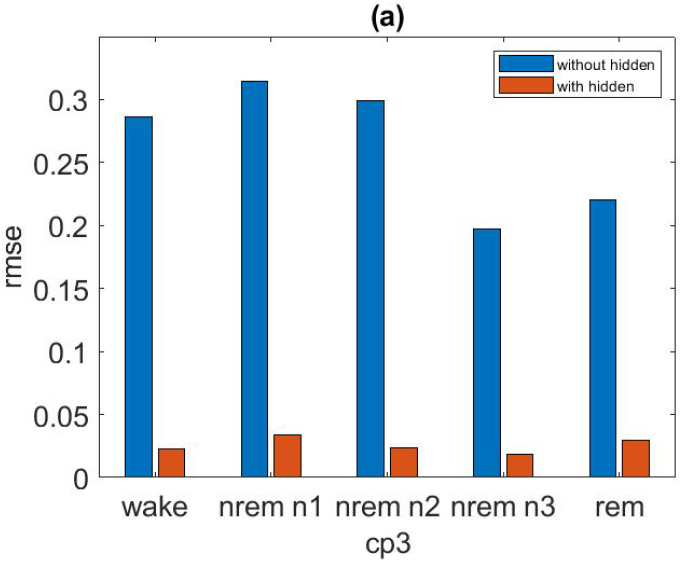
Bar plot of RMSE error comparing with ‘without hidden layer’ ‘with hidden layer’ of four different electrodes for all five sleep stages.

### Effect of hidden layer size

In the previous section, we observed enhancement of model performance after adding additional hidden layer which consists 100 sigmoidal neurons. Here we vary the number of hidden neurons in the hidden layer and compare the corresponding reconstruction RMSE (fig. 9). As expected, increasing the hidden layer size reduced RMSE over the range of hidden layer sizes shown in fig. 9. However, further increase might cause overfitting and an increase in RMSE.

**Figure 9.**
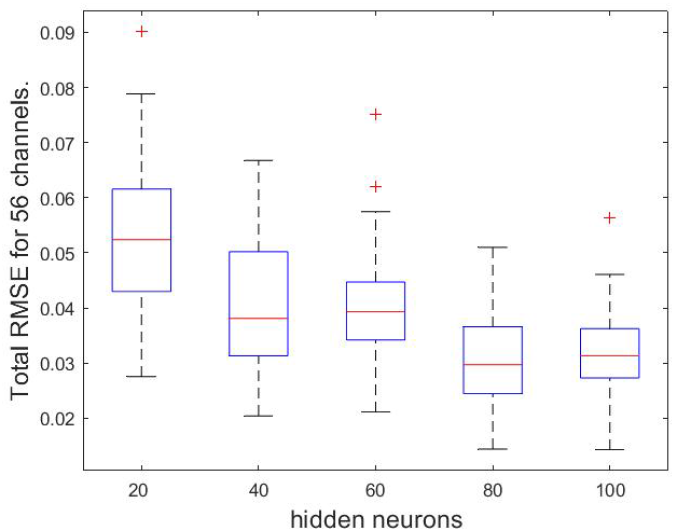
Comparing the effect of hidden layer size to reconstruct 56 EEG electrodes.

### Generation of EEG Data

Here we generate EEG data for another 30 sec using the trained parameters-intrinsic frequency of oscillators, feed-forward weights., and varying the initial phase of oscillators. Initial phases of oscillators are initialized in the interval of (0 < *ϕ*_*i*_(0) < (*απ*)) w. Where the value of *α* is 0.8. The spectrum of the generated signal is compared with next 30sec segment of training segment (which is testing data). As we observed during testing, the model is capable of generating EEG signal even beyond the training duration.

Perhaps time domain properties like spindle or k complex in NREM N2 might occur at different time periods than in the training data, but most importantly, it is retaining its overall spectral features. In fig. 9(a), the model is trained on one epoch (30 seconds in duration: 4 hrs. 12 min. 00 s. 4 hrs. 12 min. 30 s.(Table 1)) of NREM N2. During testing, the model generated signal’s spectrum is compared with the nearest next epoch (testing epoch-4hr.12min.30s-4hr.13min.00s). As we know, the sleep spindle [32] (12–15 Hz) and K complex [33] are the important features of NREM N2. We can observe the spindle activity in the model generated signal in fig.10, too. As there is a hike in power spectrum of testing data as well, similar responses we can observe in network generated signal. In conclusion, we may state that our model can generate sleep EEG signals. Generation of NREM N3 has been shown in supplementary file section 6 fig 6.

**Figure 10.**
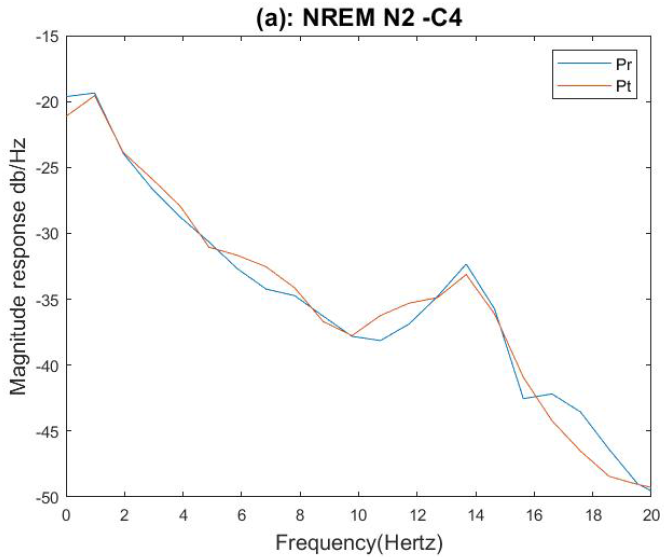
Comparison of spectral properties of network generated signal with testing signal-NREM N2 C4 channel.

**Figure 11.**
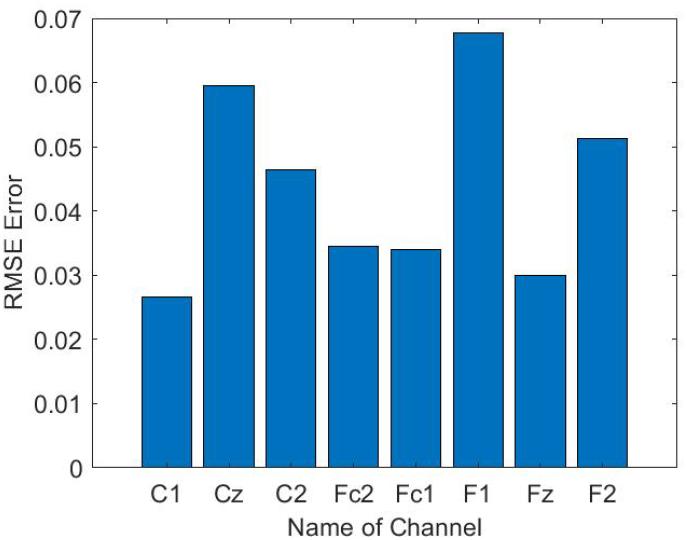
Time series of reconstructed and desired signal FC2 channel among 8 channels in a spherical shell.

**Figure 12.**
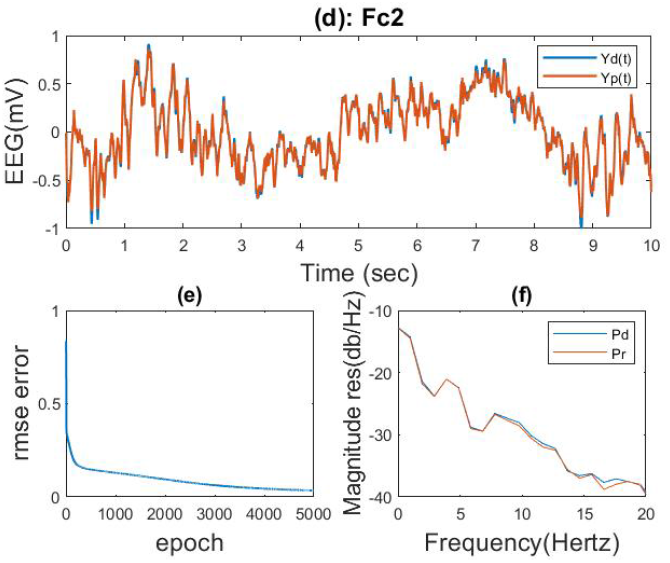
RMSE between desired and predicted signals: 8 channels and Spherical shell geometry of the cortical layer(e)-RMSE error over training, (f)-power spectrum of desired and reconstructed signal, *Y*_*d*_(*t*) =desired EEG,*Y*_*p*_(*t*)=predicted EEG,*P*_*d*_=desired EEG power.

**Table 1.** Training-testing segment of Raw EEG.

| Sleep Stage | Training Interval from raw EEG Data | Testing Interval from raw EEG Data |
| --- | --- | --- |
| NREM | 4hr:12min:00s- | 4hr:12min:30s- |
| N2 | 4hr:12min:30s | 4hr:13min:00s |
| NREM | 6hr:20min:30s- | 6hr:21min:00s- |
| N3 | 6hr:21min:00s | 6hr:21min:30s |

### Spatial distribution of oscillator

As described in Section Method. ‘spatial distribution of oscillator’, we now simulate networks in which the oscillators are spatially distributed inside a cortical layer. We consider two geometries for the cortical layer: (1) spherical shell and (2) Rectangular grid (supplementary file: Section 7)

### Spherical shell

In this study, we create a spherical surface shell (radius 85 mm obtained from EEGLab) on which the electrodes are placed. Underneath this spherical surfaceshell, we place two more spherical surfaces forming a hollow spherical shell, of inner radius (*r*_1_ = 70*mm*) and outer radius (*r*_2_ = 75 mm), within which the oscillators are distributed. Within this spherical shell we distribute 1000 Hopf oscillators. Sample locations for few channels in spherical and cartesian coordinates are shown (fig. 5 a. and 5 b.). Using a similar distance threshold (ζ_1_ = 32) defined in eqn. 15, we allocate oscillators to various electrodes (fig. 5c). The number of shared oscillators between pairwise electrodes are given in (fig. 5 d). Connectivity among oscillators is determined by the threshold value (ζ_2_ = 10).

We can see the oscillator distribution is unequal for the rectangular grid and the spherical shell. That is because of their geometrical shape. In the case of the rectangular grid, a total of 972 oscillators are distributed over three layers (see supplementary file:); the smallest shell of the rectangular grid consists of one oscillator, whereas in the spherical shell, 1000 oscillators are randomly distributed over a spherical surface. In the case of the rectangular grid, no special electrode geometry was followed –, all 8 electrodes are in the same plane, and oscillators distributed over channels are not far from each other. But in the case of spherical shell, electrode layers also having particular spherical geometrical shape, and oscillators distribution are quite random for example channel Cz is controlled by 219 number of oscillators and its neartest channel C2 is controlled by 149 oscillators. One reason might be the same fixed threshold has been used to calculate the number of oscillators belonging to each channel. Also, in comparison with a rectangular grid, a larger number of oscillators are assigned here.

## Discussion

In the present study, we model a 56-channel EEG signal with a network of oscillatory neurons. The proposed network is able to model whole-brain EEG data. The network is able to successfully extrapolate over a significant duration beyond the training duration, and is able to retain the spectral properties of the training signals over the frequency band of interest (0-20 Hz). One issue we faced during the training of a large number of EEG electrodes with a longer chunk of data was that, this model was very time consuming. For example, to train a 30-second segment of EEG data(supplimentary file) from 56 electrodes, each channel took around 2 hours of training in Intel(R) Xeon(R) CPU with 16 GB RAM type system and Matlab 2021a environment. In the present study, insertion of a hidden layer between the oscillator layer and the output layer is proven to improve reconstruction quality significantly. This is because, when there was no hidden layer, the output signal was essentially approximated by a finite set of sinusoidal signals represented by the oscillators of the input layer. But once we applied the hidden layer in our 2nd network, those sinusoid s passes through the nonlinear sigmoid functions of the hidden layer. When a single sinusoid is passes through a nonlinear sigmoid, we can get all the infinite harmonics. Hence, the hidden layer is expanding the spectrum that is available at the output layer. Thus the It was a finite number of frequencies available when we did not apply a hidden layer, suddenly explode to an infinite set of frequencies available when we we applyied a hidden layer. Also, we have successfully demonstrated spatial localization architecture for oscillator reservoirs. The performance of spatially arranged oscillators with a hidden layer is good, as demonstrated by RMSE values. However, in our spatial geometry of oscillators, we only considered locally connected regions;, but in general EEG signals from, a particular region might be influenced by local activity as well as the activity of very distant regions’ activity [21]. Also, real structural data can be implemented on this proposed model to represent a large-scale TVP kindtype of model[14]. In the future, we can easily expand our model to a real MRI-based surface where oscillators are placed according to structural-functional connectivity nodes. Thus the problem regarding the unequal distribution of oscillators in the case of “spherical shell” geometry can be eliminated. Until now we were considering one oscillator at one node (region), but in practice, many more frequency components must be present in a small node. We need to think about how we can incorporate many oscillators in a region or “network of networks” type of architecture, which can beis one possible development of step in the current work. The forward head modelling method we used in this study to explore the electrodes’ location using the BESA spherical method to extract the electrode location using EEGLab. Future efforts will be directed to developing the current model into a more realistic model of sleep dynamics. In the current modelling approach, separate networks are trained to produce EEG signals of various sleep stages and the waking stage. However, in a more authentic model of sleep, it is desirable to generate the various stages in a single model and explicitly demonstrate the transitions from one stage to the next. Ideally, such a model will show the sleep-wake cycle at a longer or diurnal time scale, and also the entire architecture of sleep substages within the 8-hour long sleep stage. The model also will permit a minimal representation of the main neural substrates of sleep regulation such as the Suprachiasmatic Nucleus (SCN), hypothalamic and thalamic nuclei and the neuromodulatory systems involved in sleep regulation, and the Reticular Activating System (RAS). In the future sleep model that we envisage, the interactions among the various subcortical circuits and neural systems produce the subrhythms of sleep, which, acting on the cortex, generate the EEG activity patterns characteristic of the relevant sleep stage.

## Acknowledgements (not compulsory)

We acknowledge the financial support from the UAY Project, funded by Govt. of India and TCS..

## Author contributions statement

The simulations were done mostly by SG with the support of DB. SG wrote the main text. VC and SV contributed to providing the key ideas, conducting initial simulations to test the hypothesis, editing the manuscript drafts, and providing insight into structure. All authors contributed to the article and approved the submitted version.

## Additional information

To include, in this order: **Accession codes** (where applicable); **Competing interests** (mandatory statement).

The corresponding author is responsible for submitting a competing interests statement on behalf of all authors of the paper. This statement must be included in the submitted article file.

