## Supplementary material for "Modeling whole brain Electroencephalogram (EEG) in a spatially organized oscillatory neural network": supplimentary

1. **The polar coordinate representation of eqn. 1(b) is:**

$$\dot{r_i} = (\mu + i\omega_i - \beta r_i{}^2)r_i + \sum_{j=1 \quad \ni j \neq i}^{N} A_{ij} r_j cos\, \omega_i \left(\frac{\emptyset_j}{\omega_j} - \frac{\emptyset_i}{\omega_i} + \frac{\theta_{ij}}{\omega_i \omega_j}\right) \qquad (1)$$

$$\dot{\emptyset_i} = \omega_i + \sum_{j=1 \quad \ni j \neq i}^{N} A_{ij} \frac{r_j}{r_i} in\, \omega_i \left(\frac{\emptyset_j}{\omega_j} - \frac{\emptyset_i}{\omega_i} + \frac{\theta_{ij}}{\omega_i \omega_j}\right) \qquad (2)$$

where $r$ and Ø are the state variables, $(\sqrt{\mu})$, ( $\mu > 0$)is the amplitude of oscillation and β is a bifurcation parameter. μ = 1, $\beta$ = −20. $A_{ij}$, is the magnitude of complex coupling coefficient, ($A_{ij} \ll 1$), $\theta_{ij}$ is angle of complex coupling coefficient, $\emptyset_i$ and $\omega_i$ and are the i[th] oscillator's phase and intrinsic frequency respectively.

2. **2[nd] stage of training**

Update rules for $K_{ij}$ and $\xi_{ij}$ are given in eqns. (3c) and (3d)

$$\Delta K_{ij} = \eta_{K_{ij}} \frac{\partial L}{\partial K_{ij}} = (-1)\eta_{K_{ij}} \sum_t (Y_{d_i}(t) - Y_{p_i}(t)) \cos(\emptyset_j(t) + \xi_{ij}) \qquad (3)$$

$$\Delta \xi_{ij} = \eta_{\emptyset_{ij}} \frac{\partial L}{\partial \emptyset_{ij}} = (-1)\eta_{\emptyset_{ij}} \sum_t \left(Y_{d_i}(t) - Y_{p_i}(t)\right)\left(-K_{ij}\, sin\,(\emptyset_j(t) + \xi_{ij})\right) \qquad (4)$$

The number of epochs for complex feedforward weight learning is 5000, and learning parameters are $\eta_{K_{ij}} = 3\text{x}10^{-5}$ , $\eta_{\emptyset_{ij}}$=$10^{-6}$.

3. **Hidden layer equation**

(learning rates: $\eta_{w_{lk}}$ =0.001; $\eta_{w_{ok}}$ =0.001).

$\dot{r_\imath} = (\mu - r_i{}^2)r_i + \sum_{j=1 \ \ni j\neq i}^{N} A_{ij} r_j \cos \omega_i \left(\frac{\emptyset_j}{\omega_j} - \frac{\emptyset_i}{\omega_i} + \frac{\theta_{ij}}{\omega_i \omega_j}\right)$ (5)

$\dot{\emptyset_\imath} = \omega_i + \sum_{j=1 \ \ni j\neq i}^{N} A_{ij} \frac{r_j}{r_i} \sin \omega_i \left(\frac{\emptyset_j}{\omega_j} - \frac{\emptyset_i}{\omega_i} + \frac{\theta_{ij}}{\omega_i \omega_j}\right)$ (6)

$$Z_r = \text{real}(\text{rarr2} * e^{i*\text{tharr2}}) \qquad (7)$$

$$Z_i = \text{imag}(\text{rarr2} * e^{i*\text{tharr2}}) \qquad (8)$$

$Z = Z_r + Z_i$ (16)

*Forward propagation:*

$W_{lk}^{f1} = W_{lk,R}^{f1} + iW_{lk,I}^{f1}$ (9)

$W_{ml}^{f2} = W_{ml,R}^{f2} + iW_{ml,I}^{f2}$ (10)

$n_l^{Hf} = \sum_k W_{lk}^{f1} Z_k = \sum_k \left(W_{lk,R}^{f1} Z_{k,R} - W_{lk,I}^{f1} Z_{k,I}\right) + i \sum_k \left(W_{lk,I}^{f1} Z_{k,R} + W_{lk,R}^{f1} Z_{k,I}\right)$ (11)

$n_l^{Hf} = n_{l,R}^{Hf} + i n_{l,I}^{Hf}$ (12)

$X_l^{Hf} = f_R^h\left(n_{l,R}^{Hf}\right) + i f_I^h\left(n_{l,I}^{Hf}\right)$ (13)

$$X_l^{Hf} = X_{l,R}^{Hf} + iX_{l,I}^{Hf} \quad (14)$$

$$n_m^o = \sum_l W_{ml}^{f2} X_l^{Hf} = \sum_l \left(W_{ml,R}^{f2} X_{l,R}^{Hf} - W_{ml,I}^{f2} X_{l,I}^{Hf}\right) + i\sum_l \left(W_{ml,I}^{f2} X_{l,R}^{Hf} + W_{ml,R}^{f2} X_{l,I}^{Hf}\right) \quad (15)$$

$$n_m^o = n_{m,R}^o + i n_{m,I}^o \quad (16)$$

$$Y_m = f_R^o\left(n_{m,R}^o\right) \quad (17)$$

*Backpropagation:*

**Loss at every time step,**

$$L(t) = \frac{1}{2}(O_m(t) - Y_m(t))^2 \quad (18)$$

$$\frac{\partial L}{\partial W_{ml,R}^{f2}} = \left(O_{m,R} - Y_{m,R}\right) f_R^{o\prime} X_{l,R}^{Hf} \quad (19)$$

$$\frac{\partial L(t)}{\partial W_{ml,I}^{f2}} = \left(O_{m,R} - Y_{m,R}\right) f_R^{o\prime} X_{m,I}^{Hf} \quad (20)$$

$$\frac{\partial L(t)}{\partial W_{lk,R}^{f1}} = (-1)\sum_m \left(O_{m,R} - Y_{m,R}\right) f_R^{o\prime} \left(W_{ml,R}^{f2} f_R^{h\prime} Z_{k,R} - W_{ml,I}^{f2} f_I^{h\prime} Z_{k,I}\right) \quad (21)$$

$$\frac{\partial L(t)}{\partial W_{lk,I}^{f1}} = \sum_m \left(O_{m,R} - Y_{m,R}\right) f_R^{o\prime} \left(W_{ml,R}^{f2} f_R^{h\prime} Z_{k,I} + W_{ml,I}^{f2} f_I^{h\prime} Z_{k,R}\right) \quad (22)$$

**Rewrite the Activation function:**

**Sigmoidal activation function:**

$$\boldsymbol{f}(x, a_k, c_k) = \frac{1}{1+\exp^{(-a_k(x-b_k))}} \quad (23)$$

Put , $a_k = 0.5, b_k = 0,$

$$2\,f(x, a_k, c_k) - 1 = \frac{1-\exp^{-0.5x}}{1+\exp^{0.5x}} = \tanh\,(x/2) \quad (24)$$

### 4. Phase locking value (PLV)

PLV measures the instantaneous phase difference, calculated using the Hilbert transform, between two narrow band signals coming from two channels. The range of the PLV value is between [0, 1], where 0 means no phase synchrony and 1 means perfectly synchronized.

PLV is calculated using

$$\boldsymbol{PLV(i,k)} = \left|\frac{1}{N}\sum_{t=1}^{N} e^{i(\Delta\emptyset)}\right| \quad \textbf{(25)}$$

For N time points, it calculates the average of N unit phasors that represent phase differences**.**

$$\boldsymbol{\Delta\emptyset = \emptyset_1 - \emptyset_2} \quad \textbf{(26)}$$

After calculating the phase, computation of PLV is done by using eqn. 4(a). Here we estimate Phase Locking Value (PLV) of experimental EEG data as well as model predicted EEG data.

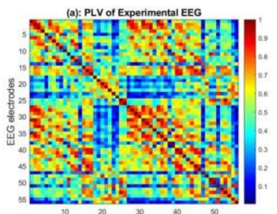

Fig 1: (a): FCM for experimental EEG Data, (b): FCM of network predicted signal using 'without hidden layer network ', (d) FCM of network predicted signal using 'with hidden layer network' .

### 5. Generation of EEG Data

$$\dot{r_i} = (\mu + i\omega_i - \beta {r_i}^2) r_i + \sum_{j=1 \quad \ni j \neq i}^{N} A_{ij} r_j cos\, \omega_i \left( \frac{\phi_j}{\omega_j} - \frac{\phi_i}{\omega_i} + \frac{\theta_{ij}}{\omega_i \omega_j} \right) \quad (27)$$

$$\dot{\phi}_i = \omega_i + \sum_{j=1 \quad \ni j \neq i}^{N} A_{ij} \frac{r_j}{r_i} in\, \omega_i \left( \frac{\phi_j}{\omega_j} - \frac{\phi_i}{\omega_i} + \frac{\theta_{ij}}{\omega_i \omega_j} \right) \quad (28)$$

### 6. Training with longer duration EEG

Using the model architecture as described in Sections II (D-F), we train the model to learn EEG signals of longer durations (30 sec) instead of 10 sec considered above. Note that, the number of oscillators in the oscillatory layer is 200. Here also we compare the reconstruction RMSE of the model predicted and the original signal for all 56 electrodes and report performance 'without hidden layer' and 'with layer model'.

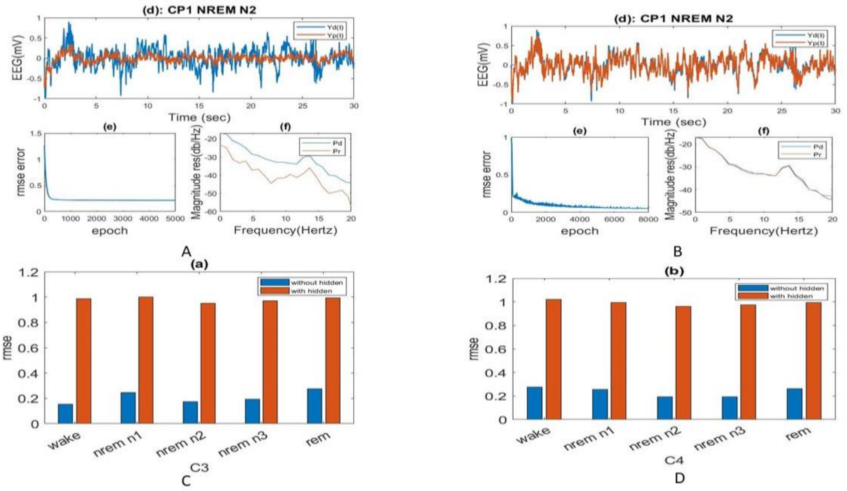

Fig 2: a) Time series reconstruction without hidden layer; b) Time series reconstruction after addition of hidden layer; c-d) bar plot of RMSE error comparing with 'without hidden layer' & 'with hidden layer' of four different electrodes for all five sleep stages.

7. **Spatial Distribution of Oscillators**

**(i) Rectangular Grid:**

Here we use 10 sec data of 8 electrodes from frontal and central lobes shown in fig 6(a-b). The signals from the 8 electrodes selected are modeled using the architecture described in section 2.8. Since the mapping between the oscillators and the electrodes is determined by the nearest neighbor criterion (eqn. 4a), it is expected that nearby electrodes are likely to share some oscillators. Training of the network proceeds as described below.

The network associated with each individual electrode - the hidden layer from which the electrode receives inputs, and the set of oscillators from which that hidden layer receives inputs – is trained separately (fig. 3). Here all 8 electrodes are selected, which form a typical rectangular grid, and the location of the electrodes in the Cartesian coordinate system is specified in Table (1). Training of the network associated with each electrode is done successively, from electrode to the next. Training begins with electrode C3 and covers all the electrode following the dotted arrows in fig. 3b.

| Channel Name | X | Y | Z |
|---|---|---|---|
| C1 | 10 | 10 | 4 |
| Cz | 20 | 10 | 4 |
| C2 | 30 | 10 | 4 |
| Fc2 | 30 | 10 | 4 |
| Fc1 | 10 | 10 | 4 |
| F1 | 10 | 10 | 4 |
| Fz | 20 | 20 | 4 |
| F2 | 30 | 20 | 4 |

Table 1: Location of 8 channel electrodes

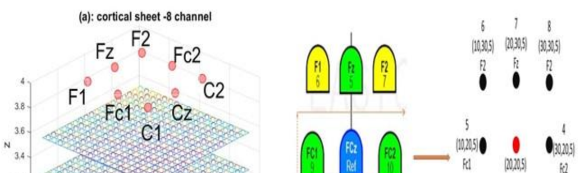


| Rectangular Grid layers | Length (mm) (X) | Breadth(mm) (Y) | Depth(mm) (Z) |
|---|---|---|---|

Fig3: a)3-layer cortical surface; b) Spatial map of Electrodes; c) Dimension of Rectangular Grid (d) The number of oscillators allocated to each channels; e) shared oscillators among the pair of channel; f) RMSE between desired and predicted signal presented for 8 channels in case of cortical layer with rectangular slab geometry.

We create a three-layer cortical surface with a rectangular geometry of dimensions (Fig. 3a and 3c). Each layer consists of 18 X 18 = 324 columns, and each column contains one oscillator. Hence totally 18 X 18 X 3 = 972 oscillators are present in the three layers. The layer of electrodes is placed on top of the 3 layered cortical surface. Next, we calculate the number of oscillators allocated to each electrode using eqn. (6a) with a distance threshold ($\xi_1 = 7.5$). All the oscillators in the cortical layer, whose distance from the individual electrode is less than ($\xi_1$), are allocated to that particular electrode. The number of oscillators allotted to various electrode are shown in Fig 3 d. Also, there is a separate hidden layer (consisting of 15 sigmoidal neurons) associated with each electrode. Note that channels that are close to each other in the electrode layer are likely to have more shared oscillators. Fig. 3 (e) provides the information about the number of shared oscillators among pairs of electrodes.

There is another threshold distance among oscillators ($\xi_2 = 1$), which determines the lateral connectivity among the oscillators using eqn. (4b). Only oscillator pairs whose mutual distance is less than $\xi_2$ are connected. The network thus designed, - with the oscillators connected among themselves over a neighborhood, and connected to the channels within a distance, with a separate hidden layer allotted to each electrode, - is trained on 8 channel EEG data.
